# Automated Annotation of Plant Gene Regions Using Supervised Machine Learning

**DOI:** 10.1101/2025.11.10.687603

**Authors:** Bruna Simões, Armando J. Pinho, Diogo Pratas

## Abstract

Rapid sequencing now enables analysis across many species, including plants, but repeat-rich, redundant plant genomes still hinder precise annotation. Precise gene calls are vital for understanding biology, comparative genomics, and discovering traits such as improved nutrition and disease resistance. We present GeAnno, a supervised machine learning method for plant gene detection. GeAnno uses an XGBoost classifier trained on curated plant annotations and a sliding-window scheme while capturing redundancy, base composition, and start/stop-codon spacing, among others. At inference, windows are scored as genic or intergenic and lightly smoothed. The output is a standard GFF3 with strand-specific genic regions. Benchmarking against ab initio predictors under matched training on 11 cassava (*Manihot esculenta*) genomes and cross-species evaluation, GeAnno achieved higher nucleotide-level precision and F1-score. For example, on cassava it reached 77.13% precision and 72.90% F1-score. Moreover, performance transfers to divergent species and parameters (window, step, smoothing, thresholds) are tunable. By improving accuracy and portability on complex plant genomes, GeAnno supports downstream functional studies and breeding, advancing food security and ecological sustainability.

## 1 Introduction

Current developments in sequencing technologies have increased exponentially the number of sequenced genomes. Next Generation Sequencing (NGS) and long-read sequencing advances have enabled the first telomere-to-telomere (T2T) assemblies of plant genomes, with the Cassava (*Manihot esculenta*) being one of the first completely sequenced plant species [6, 14].

As such, more reliable means of automated gene finding and annotation tools have gained increasing importance given this scenario, especially in the context of repeat-rich, redundant genomes, as is the case of plants [20, 7, 6, 17]. The first developed gene predictors and genome annotation tools were based mostly on HMMs (Hidden Markov Models), and trained upon another species to output its predictions (*ab initio* tools). However, these later started incorporating additional data sources such as data from Ribonucleic Acid Sequencing (RNA-seq) or other gene predictions, with comparative genomics (evidence-based tools).

Even so, the annotation of eukaryotic genomes is more challenging in the presence of variable distances between genes, alternative splicing, and higher frequency of Transposable Elements (TE) and pseudogenes [20]. Plant genomes are especially difficult to annotate due to their large DNA sequence, high number of TE and variable ploidies [22]. Most animals are diploids and contain fewer repetitive sequences comparatively to plants’ genomes. For example, using JARVIS2 and JARVIS3, the human genome compresses to about 31% of its original size, whereas plant genomes typically compress to 64%, denoting plants’ greater sequence redundancy [13, 18].

In this case, machine learning methods are considered a feasible solution to deal with these exceptions. Unsupervised learning methods have already been used to find functional elements in the human genome, and supervised learning methods, with training from manually revised annotated genomes, are also considered a solution to address the complexity of annotation [5].

As such, we present GeAnno, an approach to gene finding using supervised machine learning models. We construct several pre-trained models as means to compare with existing gene prediction and genome annotation tools, such as AUGUSTUS [19], GeneMark-ES [10], SNAP [9], GeneMark-ETP [4], GeneMark-EP+ [3] and GeMoMa [8], as well as a final model trained upon *M. esculenta* variants.

## 2 Methodology

### 2.1 Database preparation

Four different datasets were prepared to train the developed models. Three were designed to compare as closely as possible existing *ab initio* models, while the fourth, aggregating several *M. esculenta* variants, extracted from the CassavaMDB Database, was used to train the final model. Specifically, two species-specific datasets were prepared, one for *A. thaliana* and another for *O. sativa* to benchmark against the SNAP [9] and AUGUSTUS [19] pre-trained models. The third dataset encompasses data from the same species used to train the self-trained GeneMark-ES [10] *ab initio* tool, namely *A. thaliana, C. elegans* and *D. melanogaster*, extracted from the Ensembl and Ensembl Plants databases [1, 21].

Annotation and DNA files are intersected with pre-defined windows of 1500 nucleotides, separated by 50 nucleotides (step size). Bedtools’ `intersect` function [15] is used to retrieve windows within annotated gene regions, and extracting the associated sequences, labeling them as genic. All windows that do not intersect with genic regions are considered intergenic.

This logic is performed for both DNA strands, separating regions from the 5’-3’ (forward) direction and 3’-5’ (reverse) direction.

### 2.2 Feature extraction

After obtaining all window sequences and their respective strand direction, biologically relevant features are extracted. First, EMBOSS’ `getorf` [16] program is used to extract several ORFs, from which the longest found is retrieved, followed by its normalization, that is, dividing by the sequence size (in this case, 1500). This function also outputs the sequence of amino acids derived from the ORF. Therefore, this sequence is used to count the number of hydrophobic amino acids, namely, glycine, proline, phenylalanine, alanine, isoleucine, leucine, valine, tryptophan, and methionine, and subsequently, normalizing by dividing by the size of the ORF.

Another important feature is provided by the lossless genomic sequence encoder JARVIS3 [18]. JARVIS3 is used to calculate the normalized compression of the sequence as performed in previous works [12]. Sequences with higher number of low-complexity regions (LCRs) are more compressed, and can therefore be a good indicator of promoter or intergenic regions [11].

Additionally, GC content, namely the percentage of cytosine and guanine bases within a sequence, is also extracted and is associated with gene density and expression levels.

Motif distribution is also a relevant aspect, since gene expression can be associated with distances between different motifs [2]. The average distance is extracted according to

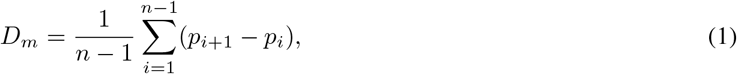

where *p*_*i*_ denotes the position of the *i*-th occurrence of motif *m*, and *n* is the number of occurrences. The median distance is given by

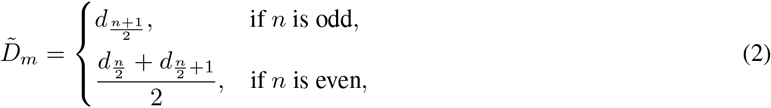

where *d*_*j*_ represents the *j*-th ordered distance.

We extracted these two distances for a set of motifs, including `A`, `C`, `T`, `G`, `ATG`, `TAA`, `TAG`, `TGA`, `TATAAA`, `CAAT`, `CGCG`, `GT`, `AG`, `AAA`, `TTT`, `GGG`, `CGG`, and `CAG`.

Finally, all frequencies of combinations of dinucleotides and tri-nucleotides present in each window were counted. These counts are then normalized by the total number of valid k-mers observed in the window, yielding relative frequencies, which are then used as features for the developed model.

### 2.3 Model training

Each developed dataset was divided into two sets, where 30% of the original data was stored for testing purposes, while 70% was used for training and validation purposes. Intergenic regions are usually larger than genic ones, presenting a class imbalance. As such, we performed undersampling to the training and validation sets. A standard scaler was also used to normalize the features, obtaining a median value of 0 and standard deviation of 1.

Four initial classifiers were considered, namely, the K-Nearest Neighbors (KNN), Multilayer Perceptron (MLP), Random Forest (RF), and eXtreme Gradient Boosting (XGBoost). Among these evaluated models, RF and XGBoost achieved the best overall performance across precision, recall, F1-score, AUCROC (Area Under Receiver Operating Characteristic Curve), and AUPRC (Area Under Precision–Recall Curve) metrics. Given the comparable accuracy of both models, XGBoost was ultimately selected due to its faster prediction time and lower memory usage. These results are presented in Supplementary Section 1.1.

Model optimization was performed through randomized hyperparameter search combined with three-fold cross-validation, allowing an efficient exploration of the parameter space while maintaining computational feasibility, choosing a randomized search approach, using 10 iterations, upon the defined search space.

Finally, in addition to the baseline configuration, models based on *A. thaliana, O. sativa*, and the GeneMark-ES training set were also subjected to Principal Component Analysis (PCA) to assess the impact of dimensionality reduction, maintaining 95% of the variance of the data.

### 2.4 Tool workflow

Building on these models, GeAnno was developed to predict genic regions from a DNA file. The workflow begins by scanning the input sequence with the user-defined window and step sizes, extracting the full set of features in a similar manner to the one defined in Feature Extraction (Subsection 2.2).

Windows were generated for both DNA strands (forward and reverse) and feature sets were extracted accordingly. The trained model then receives these features as input and outputs, for each window, the probability of belonging to a genic region. These probabilities are subsequently processed through a moving average filter, which smooths the predictions. A user-defined threshold is then applied to identify contiguous genic regions. The resulting segments are then merged per strand, and their start and end coordinates, along with strand orientation, are written to a standard GFF3 file.

### 2.5 Implementation

The methodology is implemented in a Python tool released under the MIT license (https://github.com/cobilab/GeAnno). The repository includes trained models and steps to install and use the tool. Supplementary Section 2 details its installation, parameters, and usage. The scripts to reproduce the benchmarks are released under the MIT license (https://github.com/Brums21/benchmark). Supplementary Section 3 details these steps.

## 3 Benchmark

To benchmark GeAnno, we used DNA and annotation files from four species, namely, *A. thaliana, G. raimondii, M. esculenta* and *O. sativa*. We benchmarked the performance of GeAnno with standard genome annotation and gene prediction tools such as AUGUSTUS [19], GeneMark-ES [10], GeneMark-EP+ [3], GeneMark-ETP [4], SNAP [9] and GeMoMa [8].

Accordingly, we compared the performance of ab initio tools using training datasets that most closely matched their original configurations. For AUGUSTUS, models were trained on *A. thaliana* for all testing species, except for *O. sativa*, which used its own training set. For SNAP, we conducted experiments using both *A. thaliana* and *O. sativa* training sets. For GeneMark-ES, we employed the same training species used in its native implementation, namely *A. thaliana, D. melanogaster*, and *C. elegans*. All models based on these three training sets were developed both with and without PCA. Then, we make an overall comparison for all benchmarked tools with GeAnno, using the *M. esculenta* variants trained model.

Figure 1 shows the performance of *ab initio* tools comparatively to GeAnno for different training sets. The developed tool consistently demonstrated overall better precision and F1-score values for species it was not trained upon, demonstrating a great generalization capacity for species it hasn’t seen before, even if they are taxonomically distant from the one being analyzed.

**Figure 1.**
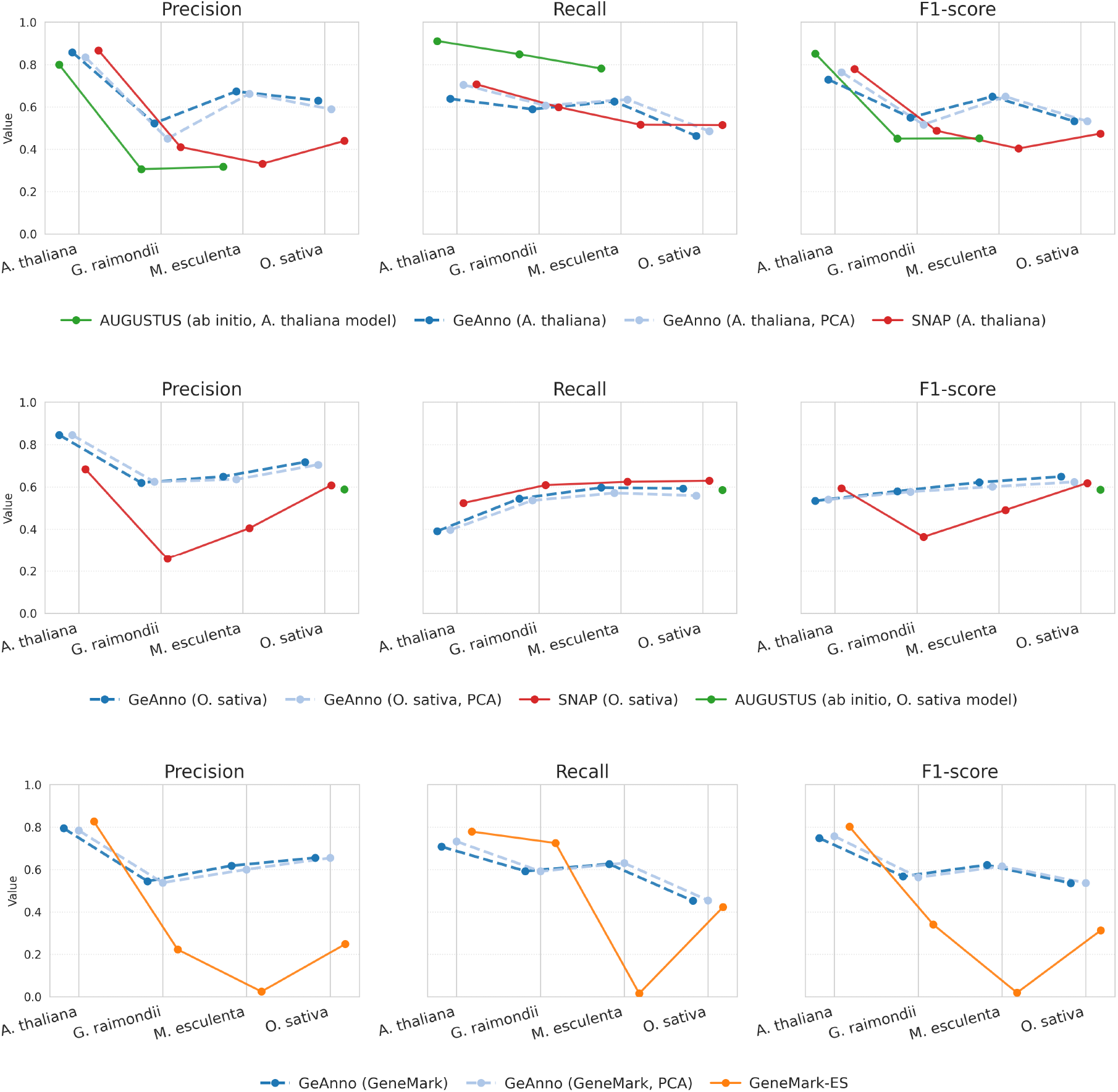
Comparison between GeAnno and the AUGUSTUS, SNAP and GeneMark-ES *ab initio* tools trained on the same species set. Each panel shows precision, recall and F1-score across benchmarked species. Dashed lines correspond to the developed tool. The first horizontal panel (from the top) shows the *A. thaliana*-trained models, the second horizontal panel shows the *O. sativa*-trained models, and the third horizontal panel shows the GeneMark-ES comparison with the same species set.

We also tested all tools across different mutation rates. We inject variants at specified frequencies to the DNA sequence to be analyzed, changing one of its nucleotides with a certain probability. For our benchmark, we performed tests at 0% (meaning the original sequence unaltered), 1%, 4% and 7% mutation rates.

Figure 2 denotes these values, at different mutation rates, where our tool presented the best precision among *ab initio* tools, and the second best comparatively to evidence-based tools, across all defined mutation rates. Recall had lower performance, but still presented the best results at higher mutation rate for all *ab initio* tools. As for F1-score, we denote the best values of GeAnno across all *ab initio* tools, and the second best for evidence-based ones, given by the second best F1-score at higher mutation rate tested (7%).

**Figure 2.**
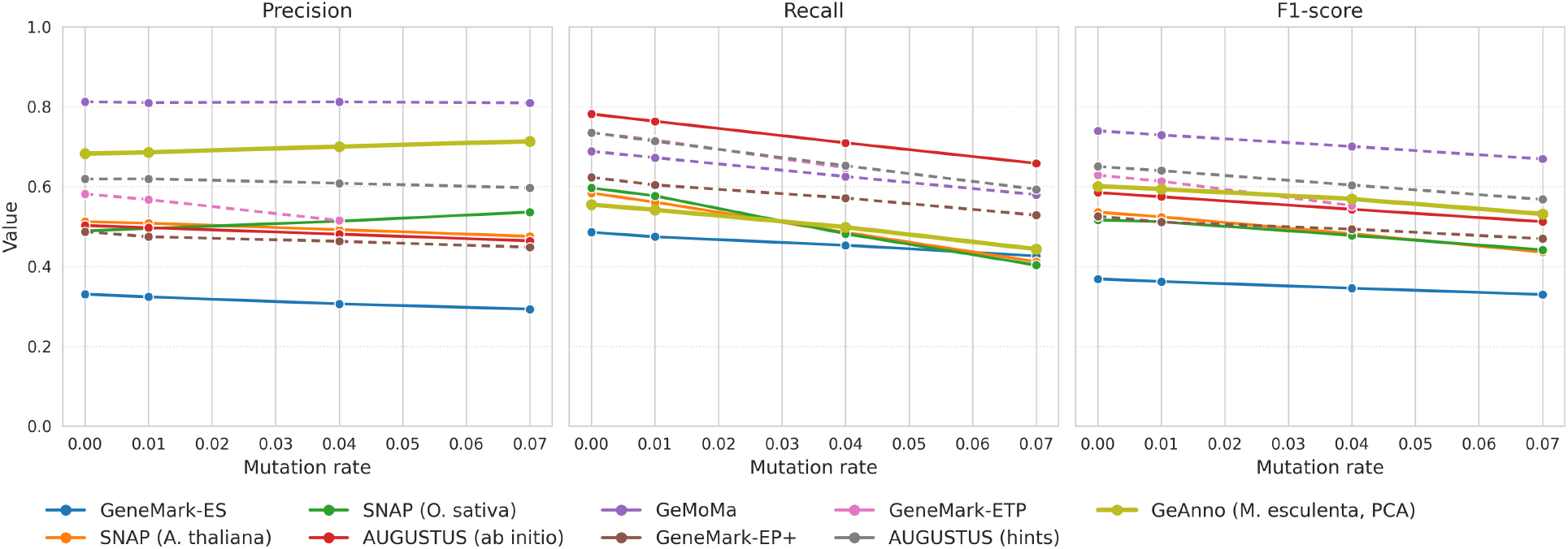
Mutation rate effect across all tools in precision, recall and F1-score metrics. We separate SNAP into the two pre-trained models used. AUGUSTUS is separated into *ab initio* and evidence-based mode. “Value” indicates the metric’s decimal value in each panel, with mutation rate also presenting decimal values. Dashed lines represent evidence-based tools, while normal lines represent *ab initio* ones. Our tool uses the final *M. esculenta* model, which is denoted as a bolder line.

Supplementary Figure S4 reports AUROC and AUPRC, computed from window-level prediction scores for GeAnno and AUGUSTUS. GeAnno attains consistently higher AUPRC, indicating greater precision among top-ranked predictions and stronger positive-class performance under class imbalance.

## 4 Conclusions

We introduced GeAnno, a Python-based tool designed for gene prediction tailored for plant genomes, using a supervised machine learning model to detect genic and intergenic regions. We tested our tool using the XGBoost classifier in four different species and compared precision, recall and F1-metrics with existing genome annotation and gene prediction tools. Under matched training on 11 cassava genomes and cross-species evaluation, GeAnno achieved higher nucleotide-level precision and F1-score than ab initio predictors. Moreover, we found that the developed tool has good generalization across different species, producing good results even for species that are taxonomically distant from the ones it was trained upon. The developed tool is available as an open-source tool, under an MIT License, ensuring accessibility and encouraging collaboration in the scientific community.

## Author contributions statement

A.P. and D.P. designed the experiment. B.S. coded the tool. B.S. executed data analysis. All authors analysed, discussed the results. B.S. and D.P. wrote the manuscript. All authors revised the manuscript.

