## Supplementary Material for "Automated Annotation of Plant Gene Regions Using Supervised Machine Learning"

B. Simões, A. J. Pinho, and D. Pratas

#### 1 Results - Supplementary material

##### 1.1 Initial Classifier Results

In an initial project phase, we analyzed several classifiers based on an initial dataset. This dataset contained data from four different species, namely *B. napus*, *H. vulgare*, *S. lycopersicum*, and *Z. mays*. These species were gathered from the Ensembl Plants website [1]. We considered only a small region from each species, and separated each region into window subsets of 1500 nucleotides, following a step of 50. Each window was classified as "genic" or "intergenic" based on the location in the annotation file: if at least 50% of the window overlapped a gene region, then that window was considered "genic"; otherwise, it's "intergenic".

The results of these classifiers are presented in Table 1.

| **Species** | **Classifier** | Precision | Recall | F1-score | AUCROC | AUPRC |
|---|---|---|---|---|---|---|
| Oversampling | KNN | 63.62 | 63.62 | 63.62 | 63.62 | 58.68 |
| | MLP | 70.21 | 70.21 | 70.21 | 70.21 | 64.25 |
| | RF | **72.03** | **71.80** | **71.72** | **71.80** | **66.19** |
| | XGB | **<u>71.71</u>** | **<u>71.55</u>** | **<u>71.50</u>** | **<u>71.55</u>** | **<u>65.85</u>** |
| Undersampling | KNN | 63.94 | 63.95 | 63.93 | 63.90 | 57.95 |
| | MLP | 69.87 | 69.25 | 69.14 | 69.47 | 65.51 |
| | RF | **71.91** | **71.64** | **71.62** | **71.76** | **67.36** |
| | XGB | **<u>71.76</u>** | **<u>71.58</u>** | **<u>71.58</u>** | **<u>71.67</u>** | **<u>67.17</u>** |

Table S1: Initial classifier metrics for four different classifiers, trained upon four different species. In this case, we considered genic windows the ones that surpass 50% overlap with a genic window. All values are percentages, with the highest being shown in a bold format, and second-highest being presented in bold and underline.

It is possible to verify that ensemble-based classifiers (RF and XGB) consistently outperformed KNN and MLP across most evaluation metrics. The differences were particularly evident in AUPRC, which is especially relevant for imbalanced datasets, as it provides a more accurate reflection of performance on the minority class.

Furthermore, undersampling generally led to slightly improved AUPRC values compared to oversampling, suggesting that reducing the size of the majority class yielded more informative decision boundaries in this context.

##### 1.2 GeAnno's configuration analysis

We ran controlled tests to verify prediction consistency across parameter choices. To observe how the tool responds to different settings, each experiment was executed with two window sizes (1000, 1500) and two step sizes (25, 50). We also verify if parameter choice is affect by different mutation rates, and if they are species-specific.

In this Table we observe consistent trade-offs, where precision tends to increase with a smaller window and larger step, but recall tends to decrease. All species achieve the best results at (1500, 50) for F1-score, except for *G. raimondii*. However, it only denotes a difference 0.01% comparatively to the highest combination)

We also denote the precision, recall and F1-score values across different thresholds, present in Figure 1.

| Metric | Species | Window and step size combination | | | |
|---|---|---|---|---|---|
| | | 1000 - 25 | 1000 - 50 | 1500 - 25 | 1500 - 50 |
| Precision | *A. thaliana* | 69.76 | **73.60** | 64.51 | 67.37 |
| | *G. raimondii* | 39.01 | **44.60** | 31.24 | 34.85 |
| | *M. esculenta* | 51.25 | **56.71** | 42.86 | 46.71 |
| | *O. sativa* | 50.77 | **55.17** | 46.63 | 49.86 |
| Recall | *A. thaliana* | 63.22 | 58.25 | **77.09** | 74.17 |
| | *G. raimondii* | 61.56 | 56.91 | **75.03** | 72.51 |
| | *M. esculenta* | 64.82 | 60.16 | **78.95** | 76.63 |
| | *O. sativa* | 54.43 | 49.81 | **66.98** | 64.37 |
| F1-score | *A. thaliana* | 62.61 | 60.92 | 67.76 | **67.97** |
| | *G. raimondii* | 42.82 | **44.16** | 41.61 | 44.15 |
| | *M. esculenta* | 53.41 | 53.93 | 53.56 | **55.81** |
| | *O. sativa* | 48.15 | 47.61 | 52.14 | **53.25** |

Table S2: GeAnno's scores across different metrics, species and combination of window sizes and steps. Columns represent the notation '<window size> - <step size>'. All values are percentages, and, for each row, bold values represent the highest, while underlined presents the lowest.

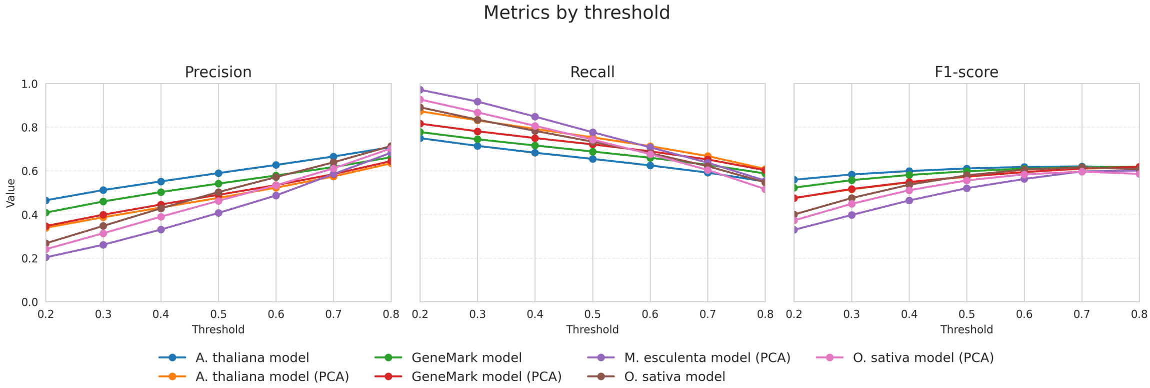


Figure S1: GeAnnos's precision, recall and F1-score metrics for different thresholds, for each developed model. Both score and threshold are shown as decimals, with "Value" indicating the metric's decimal value in each panel.

#### 1.3 GeAnno models comparison

| Metric | Species | Model | | | | | | |
|---|---|---|---|---|---|---|---|---|
| | | *A. thaliana* model | | *O. sativa* model | | GeneMark model | | *M. esculenta* model |
| | | Non-PCA | PCA | Non-PCA | PCA | Non-PCA | PCA | PCA |
| Precision | *A. thaliana* | **88.13** | 83.43 | 84.59 | 84.42 | 80.41 | 78.36 | 80.51 |
| | *G. raimondii* | 59.55 | 44.97 | 61.51 | **62.41** | 55.19 | 53.76 | 51.54 |
| | *M. esculenta* | 68.41 | 66.19 | 66.27 | 63.52 | 63.48 | 60.02 | **77.13** |
| | *O. sativa* | 67.08 | 58.90 | **72.92** | 70.46 | 65.58 | 65.44 | 63.70 |
| | **Avg.** | 70.79 | 63.37 | **71.32** | 70.20 | 66.17 | 64.40 | 68.22 |
| Recall | *A. thaliana* | 57.19 | 70.43 | 38.48 | 39.67 | 68.29 | **73.21** | 44.61 |
| | *G. raimondii* | 57.13 | 60.64 | 55.26 | 53.60 | 59.34 | 59.13 | **61.51** |
| | *M. esculenta* | 61.70 | 63.50 | 62.22 | 57.07 | 62.17 | 63.02 | **69.11** |
| | *O. sativa* | 44.12 | 48.51 | **62.59** | 55.88 | 44.99 | 45.46 | 46.62 |
| | **Avg.** | 55.04 | **60.77** | 54.64 | 51.56 | 58.70 | 60.21 | 55.46 |
| F1-score | *A. thaliana* | 69.36 | **76.38** | 52.89 | 53.97 | 73.86 | 75.70 | 57.41 |
| | *G. raimondii* | **58.31** | 51.64 | 58.22 | 57.67 | 57.19 | 56.32 | 56.09 |
| | *M. esculenta* | 64.88 | 64.82 | 64.18 | 60.12 | 62.82 | 61.48 | **72.90** |
| | *O. sativa* | 53.23 | 53.20 | **67.36** | 62.33 | 53.37 | 53.65 | 53.83 |
| | **Avg.** | 61.45 | 61.51 | 60.66 | 58.52 | **61.81** | 61.79 | 60.06 |

Table S3: GeAnno's metrics across different species for original (non-PCA) and PCA models. We used 1500 window size, 50 step size and 0% mutation rate, at 0.8 threshold. Last column corresponds to the final model used for comparisons. For each model and metric, we report the average across all benchmarked species ('Avg.' row). All values are percentages, with bold and underlined values marking the highest and lowest averages, respectively. The final delimited column corresponds to the final model used for comparison with benchmarked tools, trained upon 11 variants of the cassava species.

| **Metric** | **Mutation rate (%)** | **Model** | | | | | | |
|---|---|---|---|---|---|---|---|---|
| | | *A. thaliana* model | | *O. sativa* model | | GeneMark model | | *M. esculenta* model |
| | | Non-PCA | PCA | Non-PCA | PCA | Non-PCA | PCA | PCA |
| Precision | 0 | 70.79 | <u>63.37</u> | **71.32** | 70.20 | 66.17 | 64.40 | 68.22 |
| | 1 | 70.46 | <u>62.74</u> | **70.99** | 70.10 | 65.89 | 64.11 | 68.58 |
| | 4 | 69.64 | <u>61.21</u> | **70.95** | 70.17 | 65.52 | 64.08 | 69.98 |
| | 7 | 69.21 | <u>59.94</u> | **71.82** | 71.41 | 65.16 | 64.12 | 71.32 |
| | **Diff.** | 1.58 | **3.43** | -0.50 | -1.21 | 1.01 | 0.28 | <u>-3.10</u> |
| Recall | 0 | 55.03 | **60.77** | 54.64 | <u>51.55</u> | 58.70 | 60.20 | 55.46 |
| | 1 | 54.51 | **60.66** | 53.36 | <u>51.03</u> | 58.20 | 59.57 | 54.15 |
| | 4 | 52.80 | **60.32** | 49.92 | <u>48.95</u> | 56.58 | 57.52 | 49.82 |
| | 7 | 50.39 | **59.23** | 44.75 | 45.85 | 53.68 | 54.19 | <u>44.31</u> |
| | **Diff.** | 4.64 | <u>1.54</u> | 9.89 | 5.70 | 5.02 | 6.01 | **11.15** |
| F1-score | 0 | 61.45 | 61.51 | 60.66 | <u>58.52</u> | **61.81** | 61.79 | 60.06 |
| | 1 | 60.94 | 61.08 | 59.64 | <u>58.09</u> | **61.37** | 61.29 | 59.35 |
| | 4 | 59.38 | 59.97 | 57.09 | <u>56.59</u> | **60.17** | 60.06 | 56.87 |
| | 7 | 57.46 | **58.52** | 53.43 | 54.62 | 58.14 | 58.09 | <u>53.13</u> |
| | **Diff.** | 3.99 | <u>2.99</u> | **7.23** | 3.90 | 3.67 | 3.70 | 6.93 |

Table S4: GeAnno's metrics across different mutation rates for original (non-PCA) and PCA models. We used window size 1500, step size 50, and threshold 0.8, and averaged results across all benchmarked species. Last column corresponds to the final model used for comparisons. The 'Diff.' row reports the signed difference between metrics at 0% and 7% mutation rate for each model. Negative values indicate the metric is higher at 7% than at 0%. All values are percentages, with bold and underlined values marking the largest and smallest absolute differences, respectively.

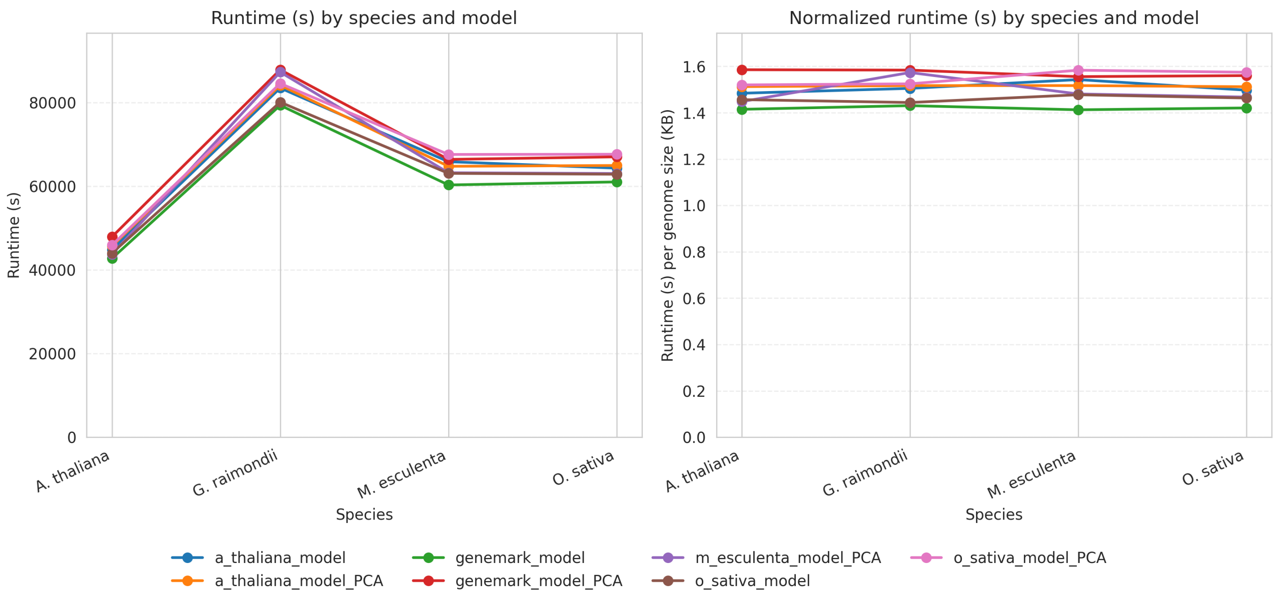


Figure S2: GeAnno's runtimes per benchmarked species. Figure on the left shows the averaged values of runtime, in seconds, per model, while the figure on the right shows the normalized average values of runtime by dividing the corresponding genome size. Legend denotes the model, and whether it used PCA or not.

#### 1.4 Benchmark results

##### 1.4.1 Comparison with *ab initio* tools

| Metric | Species | AUGUSTUS | SNAP | | GeneMark-ES | GeAnno |
|---|---|---|---|---|---|---|
| | | | *A.thaliana** | *O. sativa** | | |
| **Precision** | *A. thaliana* | 79.92 | **86.68** | 68.35 | 82.61 | 80.51 |
| | *G. raimondii* | 30.59 | 41.06 | 25.93 | 22.26 | **51.54** |
| | *M. esculenta* | 31.72 | 33.16 | 40.43 | 2.42 | **77.13** |
| | *O. sativa* | 58.81 | 43.93 | 60.78 | 24.83 | **63.70** |
| **Recall** | *A. thaliana* | **91.10** | 70.62 | 52.38 | 77.86 | 44.61 |
| | *G. raimondii* | **84.88** | 59.89 | 60.82 | 72.46 | 61.51 |
| | *M. esculenta* | **78.12** | 51.55 | 62.37 | 1.54 | 69.11 |
| | *O. sativa* | 58.49 | 51.39 | **62.91** | 42.30 | 46.62 |
| **F1-score** | *A. thaliana* | **85.14** | 77.83 | 59.31 | 80.16 | 57.41 |
| | *G. raimondii* | 44.97 | 48.72 | 36.36 | 34.05 | **56.09** |
| | *M. esculenta* | 45.12 | 40.35 | 49.06 | 1.88 | **72.90** |
| | *O. sativa* | 58.65 | 47.37 | **61.83** | 31.29 | 53.84 |

Table S5: Comparison of *ab initio* tools (AUGUSTUS [2], in its *ab initio* mode, SNAP [3], with different training species models, GeneMark-ES [4]) and GeAnno across benchmarked species and metrics. Under SNAP, we present the results for each of the models used, marked with the '*' symbol. For AUGUSTUS [2], all species use the *A. thaliana* model, except for the *O. sativa* species, which uses the *O. sativa* model. For GeAnno, we use the final *M. esculenta* variants model, with 1500 window size and 50 step size, with a 0.8 threshold. All values are percentages, with bold and underlined text representing highest and lowest values, respectively.

##### 1.4.2 Comparison with evidence-based tools

We compare GeAnno with tools that require extrinsic evidence, such as GeMoMa, GeneMark-EP+, GeneMark-ETP and AUGUSTUS. GeneMark-EP+, GeneMark-ETP and AUGUSTUS take as evidence a set of hints to support their predictions, while GeMoMa uses the annotation and DNA file of a closely related species. We followed a similar approach to the experiments performed in [5] and divided the experiments into different sets of evidence used for each evidence-based tool, namely, sets with species/hints from the same genus, same order or no close relation between the testing species.

We use the following configuration shown in Tables 6 and 7 for these tools.

Figure 3 denotes the comparison between GeAnno and the evidence-based benchmarked tools, taking into consideration its different evidence categories.

Table 8 summarizes each evidence-based tool per species, averaging scores across the available protein-hint categories so each hint type contributes equal weight.

##### 1.4.3 Comparison between AUGUSTUS and GeAnno for AUCROC and AUPRC metrics

| | | | | | |
|---|---|---|---|---|---|
| **Same genus** | **Testing** | *A. thaliana* | *G. raimondii* | *M. esculenta* | *O. sativa* |
| | **Training** | *A. lyrata*<br>*A. halleri* | — | — | *O. nivara*<br>*O. barthii* |
| **Same order** | **Testing** | *A. thaliana* | *G. raimondii* | *M. esculenta* | *O. sativa* |
| | **Training** | *E. salsugineum*<br>*B. napus* | *C. capsularis*<br>*T. cacao* | — | *Z. mays*<br>*H. vulgare* |
| **No close relation** | **Testing** | *A. thaliana* | *G. raimondii* | *M. esculenta* | *O. sativa* |
| | **Training** | *Z. mays*<br>*H. vulgare* | *Z. mays*<br>*H. vulgare* | *C. capsularis*<br>*T. cacao* | *E. salsugineum*<br>*B. napus* |

Table S6: Protein hint setup according to each species. The protein files, in FASTA format, from the two different species were combined to a singular FASTA file, which in turn was provided as the required input. Cells with the '—' notation refer to the empty sets, that is, were not considered for benchmark due to missing protein files corresponding to the respective sets in the referenced plant database. The 'Training' row corresponds to training species, while 'Testing' corresponds to testing species. 'Same genus', 'same order' and 'no close relation' correspond to same genus hints, same order hints and no close relation hints, respectively.

| | | | | | |
|---|---|---|---|---|---|
| **Same genus** | **Testing** | *A. thaliana* | *G. raimondii* | *M. esculenta* | *O. sativa* |
| | **Training** | *A. lyrata* | — | — | *O. nivara* |
| **Same order** | **Testing** | *A. thaliana* | *G. raimondii* | *M. esculenta* | *O. sativa* |
| | **Training** | *B. napus* | *C. capsularis* | — | *Z. mays* |
| **No close relation** | **Testing** | *A. thaliana* | *G. raimondii* | *M. esculenta* | *O. sativa* |
| | **Training** | *C. capsularis* | *A. lyrata* | *A. lyrata* | *A. lyrata* |

Table S7: Reference [6] species' setup used by the GeMoMa [6] tool. Cells with the '—' notation refer to the empty sets, that is, were not considered for benchmark due to missing annotation/DNA files corresponding to the respective sets in the referenced plant database. The 'Training' row corresponds to training species, while 'Testing' corresponds to testing species. 'Same genus', 'same order' and 'no close relation' correspond to same genus hints, same order hints and no close relation hints, respectively.

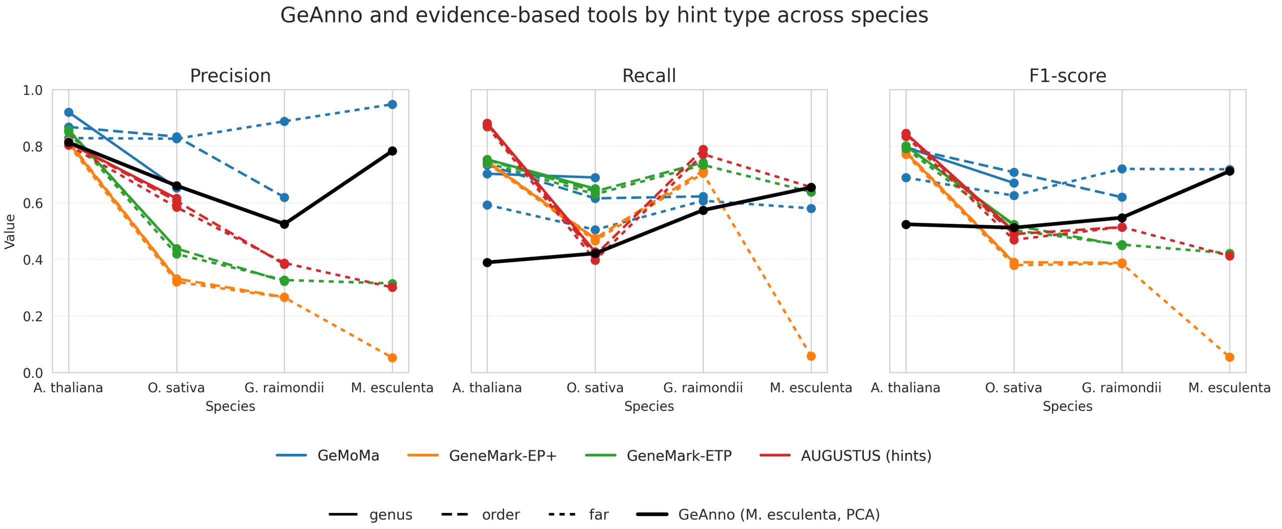


Figure S3: Comparison between evidence-based tools across different hint types and GeAnno. "Value" indicates the metric's decimal value in each panel, with each tool being represented with a different color. Dashing styles represent different hint groups.

| Metric | Species | Tools | | | | |
|---|---|---|---|---|---|---|
| | | AUGUSTUS | GeMoMa | GeneMark-EP+ | GeneMark-ETP | GeAnno |
| **Precision** | *A. thaliana* | 81.55 | **87.39** | 83.70 | 86.58 | 80.51 |
| | *G. raimondii* | 37.71 | **75.30** | 27.36 | 33.15 | 51.54 |
| | *M. esculenta* | 32.30 | **95.84** | 5.94 | 34.38 | 77.13 |
| | *O. sativa* | 61.83 | **77.16** | 34.82 | 48.50 | 63.70 |
| **Recall** | *A. thaliana* | **90.18** | 72.44 | 77.35 | 76.53 | 44.61 |
| | *G. raimondii* | **84.40** | 67.41 | 75.75 | 76.04 | 61.51 |
| | *M. esculenta* | **73.49** | 64.56 | 7.62 | 71.34 | 69.11 |
| | *O. sativa* | 50.61 | 64.16 | 52.46 | **69.33** | 46.62 |
| **F1-score** | *A. thaliana* | **85.65** | 79.18 | 80.40 | 81.25 | 57.41 |
| | *G. raimondii* | 52.12 | **70.50** | 40.20 | 46.17 | 56.09 |
| | *M. esculenta* | 44.88 | **77.15** | 6.68 | 46.40 | 72.90 |
| | *O. sativa* | 55.66 | **69.26** | 41.86 | 57.07 | 53.83 |

Table S8: Comparison of evidence-based tools AUGUSTUS [2] (using protein evidence only), GeMoMa [6], GeneMark-EP+ [7], GeneMark-ETP [5] and GeAnno across species and metrics. For each species, scores are averaged across all available protein hint categories at 0% mutation. For GeAnno, we evaluate the *M. esculenta* variants model, with 1500 window size and 50 step size. All values are percentages, and, for each row, there is an underlined and bold formatted value corresponding to the lowest and greatest value in that row, respectively.

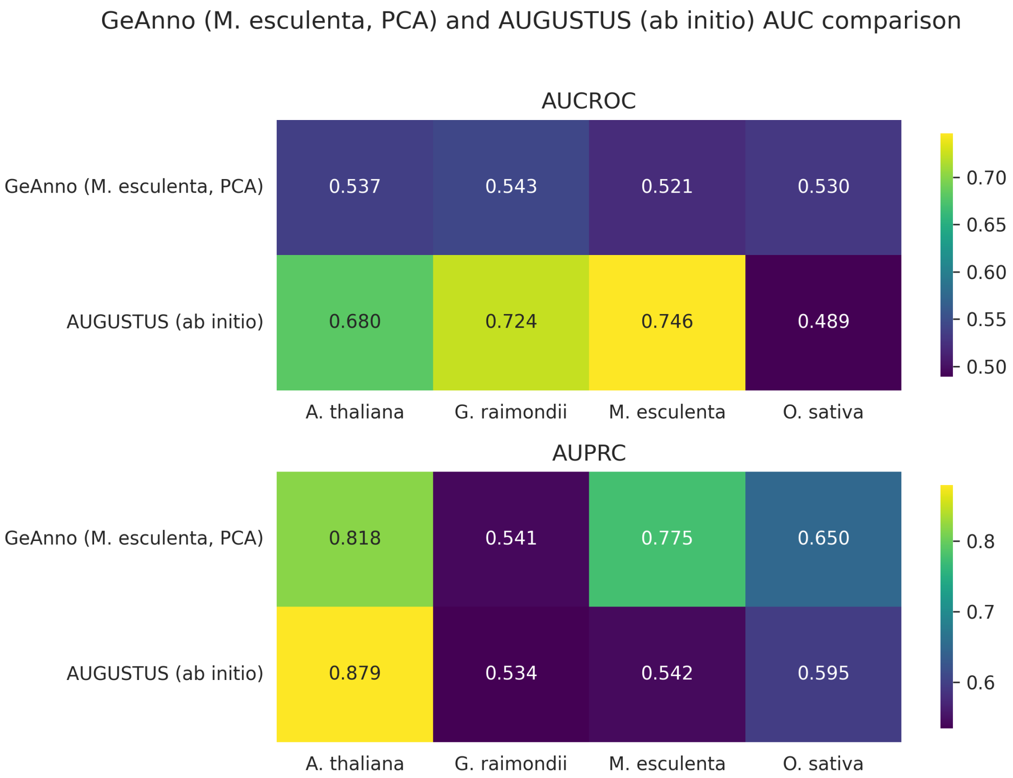


Figure S4: Comparison of AUCROC and AUPRC metrics between the developed method (GeAnno) and AUGUSTUS (*ab initio* mode). All values are shown as decimals.

### 2 GeAnno tool

#### 2.1 Installation

GeAnno can be installed using

```
git clone https://github.com/Brums21/GeAnno.git
cd GeAnno

echo "export PLANT_DIR=$(pwd)" >> env_geanno.sh
source env_geanno.sh

chmod +x ./setup.sh
./setup.sh -t

source .venv/bin/activate
```

#### 2.2 Usage

To use GeAnno on a given sequence, use

```
python3 src/geanno.py -d sequence.fa -m xgboost.pkl
```

To list the available options, use

```
python3 src/geanno.py -h
```

This command will output the following message

```
This command will output the following message:

usage: geanno.py [-h] [-d DNA_FILE] -m MODEL [-f FIGURE_FOLDER] [-o OUTPUT_FOLDER] [-ff FEATURES_FOLDER]
                 [-c CSV_FEATURES_FILE_PATH] [-p SAVE_PLOT] [-s STEP] [-t THRESHOLDS] [-w WINDOW_SIZE]
                 [-ma MOVING_AVERAGE]

options:
  -h, --help            show this help message and exit
  -d DNA_FILE, --dna_file DNA_FILE
                        FASTA DNA file (required if --csv_features_file_path not provided)
  -m MODEL, --model MODEL
  -f FIGURE_FOLDER, --figure_folder FIGURE_FOLDER
  -o OUTPUT_FOLDER, --output_folder OUTPUT_FOLDER
  -ff FEATURES_FOLDER, --features_folder FEATURES_FOLDER
  -c CSV_FEATURES_FILE_PATH, --csv_features_file_path CSV_FEATURES_FILE_PATH
  -p SAVE_PLOT, --save_plot SAVE_PLOT
  -s STEP, --step STEP
  -t THRESHOLDS, --thresholds THRESHOLDS
  -w WINDOW_SIZE, --window_size WINDOW_SIZE
  -ma MOVING_AVERAGE, --moving_average MOVING_AVERAGE
```

#### 2.3 Training new models

We provide users with all mentioned trained classifiers, present in the `models/` directory.

However, the user can train their own models providing as input DNA sequences and their respective annotation.

To generate the dataset with calculated features, use

```
chmod +x train_models/generate_dataset_model.sh
train_models/generate_dataset_model.sh -f <FASTA_DIR> -d <GFF_DIR> -o dataset.csv
```

`FASTA_DIR` represent the directory where the DNA files are located and `GFF_DIR` corresponds to the directory where the corresponding annotation files are located. The files present in both directories must have the same nomenclature, meaning that the files `x.fa` and `y.fa` must have the corresponding `x.gff3` and `y.gff3` annotation files, respectively.

To obtain the options for this script, use

```
train_models/generate_dataset_model.sh -h
```

This command will output the following content

```
Generate dataset for genome annotation.

Syntax: train_models/generate_dataset_model.sh [-h] -f FASTA_DIR -d ANNOTATION_DIR [-w
    WINDOW_SIZE] [-s STEP] [-o OUTPUT]

options:
-h   Print this help and exit
-f   Path to folder containing FASTA files (mandatory)
-d   Path to folder containing GFF3 annotation files (mandatory)
-w   Window size (default: 1500)
-s   Step size (default: 50)
-o   Output file name (default: dataset_<WINDOW>_<STEP>.csv)
```

To train the models with the developed dataset, use

```
python3 train_models/train_model.py -d dataset.csv -m RF XGBoost
```

To obtain the options for this program, use

```
python3 train_models/train_model.py -h
```

This command will output the following content

```
usage: train_model.py [-h] -d DATASET_FILE [-m {RF,XGBoost} [{RF,XGBoost} ...]] [-o
    OUTPUT_FILES [OUTPUT_FILES ...]]
                      [--output_dir OUTPUT_DIR] [-ov] [--cv CV] [--n_jobs N_JOBS] [--
    max_workers MAX_WORKERS]
                      [--n_iter N_ITER]

options:
  -h, --help            show this help message and exit
  -d DATASET_FILE, --dataset_file DATASET_FILE
  -m {RF,XGBoost} [{RF,XGBoost} ...], --models {RF,XGBoost} [{RF,XGBoost} ...]
                        Which models to train: RF, XGBoost, or both
  -o OUTPUT_FILES [OUTPUT_FILES ...], --output_files OUTPUT_FILES [OUTPUT_FILES ...]
                        Optional .pkl filenames (one per model). If omitted, defaults are auto
    -generated.
  --output_dir OUTPUT_DIR
                        Directory where models will be saved (default: models/trained_models)
  -ov, --oversampling   Use oversampling (default: undersampling)
  --cv CV               Number of CV folds for RandomizedSearchCV
  --n_jobs N_JOBS       Number of parallel jobs for training
  --max_workers MAX_WORKERS
                        Max workers for parallel model training
  --n_iter N_ITER       Number of iterations for RandomizedSearchCV
```

### 3 Reproducibility

#### 3.1 Reproducing the Benchmark

In order to download the training sets, reference species and hints and tools, as well as their dependencies, use

```
echo "export BENCHMARK_DIR=$(pwd)" >> env.sh
source env.sh
bash ./setup.sh
```

In the `benchmarking_scripts/` folder of the JARVIS3 repository, please type

```
bash ./run_all.sh
```

The output annotations are present in the `results/tools/` folder. To compile these annotations, use

```
./metrics/extract_all_values.sh
```

After executing this script, to plot the figures and CSV with relevant metrics, use

```
cd ${BENCHMARK_DIR}/plots

python3 -m venv .venv
source .venv/bin/activate

pip install -r requirements.txt

python3 generate_all_graphics.py \
        --csv_dir ${BENCHMARK_DIR}/results/compiled/ \
        --fig_dir <output_path_to_place_figures> \
        --results_geanno ${BENCHMARK_DIR}/results/GeAnno \
        --geanno_auc_csv ${BENCHMARK_DIR}/results/GeAnno/auc_csv/geanno_auc.csv
```

#### 4 Computing environment

In our study, the benchmark was executed on a computer running Ubuntu 22.04.4 LTS (Jammy Jellyfish) with kernel version 6.8.0-60-generic, equipped with four Intel® Xeon® E7320 processors (16 cores in total) operating at 2.13 GHz and 256 GB of RAM, using the x86_64 architecture.

[7] Tomáš Brůna, Alexandre Lomsadze, and Mark Borodovsky. GeneMark-EP+: Eukaryotic gene prediction with self-training in the space of genes and proteins. *NAR Genomics and Bioinformatics*, 2, 6 2020.
